# The perception of realism is correlated with the concept of physical gamut

**DOI:** 10.64898/2026.04.20.719688

**Authors:** Killian Duay, Takehiro Nagai

## Abstract

Realism and naturalness remain unresolved questions in vision science. This study investigates whether the physical gamut correlates with realism judgements. We conducted psychophysical experiments where observers judged the realism of natural scenes with target regions manipulated across the CIE 1931 color space. Results initially showed a moderate-to-strong correlation between judgements and a theoretical physical gamut derived from optimal colors. Further analysis revealed that the most detrimental points were in the saturated green region of the CIE 1931 *xy* chromaticity diagram; removing them yielded a very strong correlation. To explain this discrepancy, we modeled a real-world physical gamut based on USGS and ECOSTRESS spectral libraries. The analysis revealed that the detrimental green chromaticities might be non-existent in the real-world. Since physical gamut theory posits that the visual system constructs internal references through empirical observation of the world, the absence of these colors in nature might be a plausible explanation to the theoretical model’s failure. Ultimately, the real-world gamut exhibited an even stronger correlation with judgements, supporting our hypothesis while suggesting that the theoretical model may not be the optimal approximation of the actual physical gamut. These findings contribute to discussions on perceptual realism and offer a framework for enhancing rendering technologies.

## Introduction

The perception of realism and naturalness, along with their underlying mechanisms, remains a highly active yet unresolved field of research. Elucidating how humans perceive these attributes is fundamental not only for advancing basic vision science but also for developing practical engineering applications.

Previous studies have demonstrated that specific manipulations of image colors, such as rigid rotations in CIELAB or CIELUV space to shift hue while preserving lightness and saturation or altering saturation while maintaining lightness and hue, significantly reduce both image preference and perceived naturalness compared to original versions^1,2^. These findings suggest an interdependency between lightness, saturation, and chromaticity in shaping the perception of naturalness. However, the precise nature of these interactions and the specific perceptual mechanisms upon which they rely have not yet been fully characterized. Nakano et al.^3^ proposed that the interaction between luminance and color saturation is central to the perception of naturalness. Their findings indicate that naturalness decreases when saturation is manipulated while luminance remains constant, or conversely, when luminance is altered while chromaticity is preserved. They further demonstrated that this interaction is governed by color appearance modes: specific shifts in the relationship between luminance and saturation can cause a color to transition from “surface color mode,” where the color is perceived as a reflective surface^4^, to “aperture color mode,” where it is perceived as self-luminous^4^. They explained that if a manipulation causes an object to appear self-luminous (aperture mode) when it is contextually unlikely to emit light, the scene is perceived as unnatural.

Consequently, predicting the threshold at which an object transitions from a reflective surface to a self-luminous source could provide a reliable indicator of image naturalness and realism. However, predicting these self-luminosity thresholds remains a challenge. While foundational studies have established initial frameworks for understanding these boundaries, these earlier models are insufficient to fully explain or accurately predict them^5–12^. More recently, studies have introduced a novel approach that predicts self-luminosity thresholds with high precision^13,14^. These findings demonstrate that luminosity thresholds are strongly correlated with the physical gamut, which can serve as a reliable predictor for self-luminosity. The concept of physical gamut represents the set of all physically possible colors in a scene under a specific illuminant; more precisely, it defines the maximum luminance a given chromaticity can attain under a particular lighting. In practical terms, modeling the physical gamut by accounting for every reflectance present in the real-world is an intractable task. Consequently, these studies rely on the concept of “optimal colors”^13,15–18^ to approximate this gamut. An optimal color is a theoretical construct derived from the spectral integration of an illuminant with a reflectance spectrum that reflects either 0% or 100% of light at any given wavelength. Since no real-world object can reflect more than 100% of incident light, the optimal colors define the theoretical maximum luminance for any chromaticity. By modeling all combinations of reflectance spectra satisfying this condition and integrating them with a specific illuminant, one obtains the physical gamut under that particular lighting condition (further technical details are provided in the Methods section). As optimal colors are a theoretical concept, the physical gamut derived from them is likewise theoretical; throughout this article, we refer to it as *theoretical physical gamut*. As aforementioned, the utility of the theoretical physical gamut has been demonstrated by Morimoto et al.^13^ and Duay and Nagai^14^, who showed a robust correlation between the gamut and the luminosity thresholds. Their works highlight the model’s ability to accurately predict the transition between surface color and self-luminosity appearance modes. It is hypothesized that the human visual system has indirectly constructed an internal reference of physical gamuts through empirical observation of the real-world across diverse lighting conditions over evolutionary timescales. According to this theory, the visual system first estimates the scene illuminant and then retrieves the corresponding physical gamut from its internal references to infer perceptual judgements. While the concept of physical gamut was previously explored by Speigle and Brainard^11^ through empirical modeling based on Munsell color chips, the theoretical model derived from optimal colors currently appears to be the most effective method for approximating real-world physical limits and predicting self-luminosity.

Consequently, as we are now able to predict self-luminosity thresholds with high precision utilizing this new approach, it would be of interest to apply it to the explanation proposed by Nakano et al.^3^ regarding the interaction between naturalness and self-luminosity. Specifically, it would be worthwhile to evaluate whether a correlation could be observed between the theoretical physical gamut and naturalness, and to determine if this gamut can predict the perception of realism. The present study was specifically designed to address this objective: our work aimed to investigate whether the theoretical physical gamut correlates with perceived realism and to evaluate its efficacy as a potential predictor of realism judgements. To this end, we conducted a psychophysical experiment in which observers assessed the perceived realism of various natural scenes. The stimuli consisted of four images in which the chromaticity and luminance of a target region were systematically changed throughout the experimental trials. The full extent of the CIE 1931 *xyY* color space has been evaluated, within the constraints of the monitor. This strategy allowed us to compare observers’ judgements against the theoretical physical gamut. Our results demonstrate a moderate-to-strong correlation between observers’ judgements and the theoretical physical gamut. Further analysis revealed that the colors located in the highly saturated green region of the CIE 1931 *xy* chromaticity diagram were primarily responsible for the degradation of this correlation. When these specific chromaticities were excluded from the correlation analysis, a very strong correlation emerged. A plausible explanation for the negative impact of the green region would be that such chromaticities do not occur in the real-world. As the physical gamut hypothesis posits that the human visual system constructs an internal reference through the empirical observation of the world over biological evolution, the absence of these colors in nature would prevent the visual system from learning them. Consequently, while a theoretical model based on ideal definitions includes these colors, it could not accurately predict their perceived realism in natural scenes. This potential explanation is supported by a secondary analysis in which we constructed a *real-world physical gamut* using measurements of reflectances from the two extensive USGS^19^ and ECOSTRESS^20,21^ spectral libraries. Our analysis showed that the real-world physical gamut contains no samples within the problematic green chromaticity region, suggesting that these colors might indeed be absent from the real-world. A final analysis demonstrated a very strong correlation between the real-world physical gamut and observers’ judgements. Notably, the real-world model outperformed the theoretical physical gamut when evaluated against the same experimental data. These findings further reinforce the hypothesis that the human visual system relies on an internal reference shaped by the empirical observation of the world. However, it should be noted that while these explanations appear plausible, they cannot be completely verified or confirmed. Indeed, they rely on strong assumptions and a body of data that, although extensive, cannot be regarded as exhaustive. In addition, a replication of the experiment under a different illuminant indicated that the findings are not restricted to a neutral illuminant. These results appear generalizable to other illuminants, thereby reinforcing the robustness of the proposed theoretical framework.

These results offer novel insights into the perception of realism. They can contribute to the broader discourse on the perception of naturalness or, indirectly, color preference. Beyond theoretical contributions, this work could be applied to help solve concrete problems encountered in the field of engineering, such as realistic graphical rendering of virtual objects or virtual worlds. More specifically, it could be beneficial for Augmented Reality (AR) or Mixed Reality (MR). Indeed, these technologies aim to integrate virtual objects into the real world with the ultimate goal of creating a seamless illusion where it becomes impossible to distinguish the virtual from the real. A primary challenge in achieving this objective is ensuring that the integration of virtual objects faithfully respects lighting constraints: specifically, plausible colors and brightness within a given real-world environment. A bright white or light blue virtual object may indeed appear unrealistic within a sunset scene characterized by intense reddish lighting, as it would violate the physically realizable gamut constraints under such illumination. The physical gamut model (including both the theoretical and real-world physical gamuts) discussed in the present study helps define the boundaries that virtual objects must adhere to in order to be perceived as realistic within Extended Reality (XR) experiences.

To ensure terminological clarity before going further, *realism* and *naturalness* are defined in the present study as inter-changeable concepts referring to the plausibility of objects or scenes to be observed in the real-world. This usage is consistent with the literature cited in this introduction. While some studies define *naturalness* in opposition to *man-made* (e.g., urban areas)^22,23^, this definition differs from the one employed in our work or in the literature introduced earlier.

## Results

The objective of this study was to evaluate the hypothesis that a correlation exists between observers’ realism judgements of natural scenes and the theoretical physical gamut; such a correlation would suggest that the gamut could serve as a predictor for the perception of realism. The theoretical physical gamut is a model representing the range of physically realizable colors under a given illuminant. We designate this model as theoretical within the current study as it is derived from the theoretical construct of optimal colors rather than empirical observations (see Methods). Recent studies have demonstrated that the theoretical physical gamut appears to be the most robust model for predicting self-luminosity thresholds to date (the critical point at which the perception of an object shifts from a light-reflecting surface to a light-emitting source)^4,13,14^. In a separate context, research has indicated that the perception of naturalness seems to be contingent upon the perception of self-luminosity. Consequently, we hypothesized that theoretical physical gamut may correlate with the perception of realism and could thus serve as a reliable predictor. To evaluate the hypothesis, a psychophysical experiment was performed wherein observers were asked to assess the realism of natural scenes presented in images. The evaluated scenes comprised four images in which the chromaticity and luminance of a target flower were systematically varied throughout experimental trials. The trials covered the entirety of the CIE 1931 *xyY* color space within the constraints of our monitor, thus enabling a subsequent comparison of observers’ judgements against the theoretical physical gamut.

Figure 1 and its interactive three-dimensional (3D) counterpart Supplementary Fig. S1 online provide a visual representation of the results, plotting observers’ judgements alongside the theoretical physical gamut within the CIE 1931 *xyY* color space. For an enhanced visualization experience, we recommend consulting the interactive version online. Observers’ judgements were smoothed, and the perceptual boundary between realistic and non-realistic stimuli was derived for each chromaticity. This processing resulted in a bell-shaped surface that delineates the limits of perceptual realism within the CIE 1931 *xyY* color space under the experimental illuminant (see Methods). In contrast, the theoretical physical gamut defines the upper limit of physically achievable luminance for each chromaticity within the same space.

**Figure 1.**
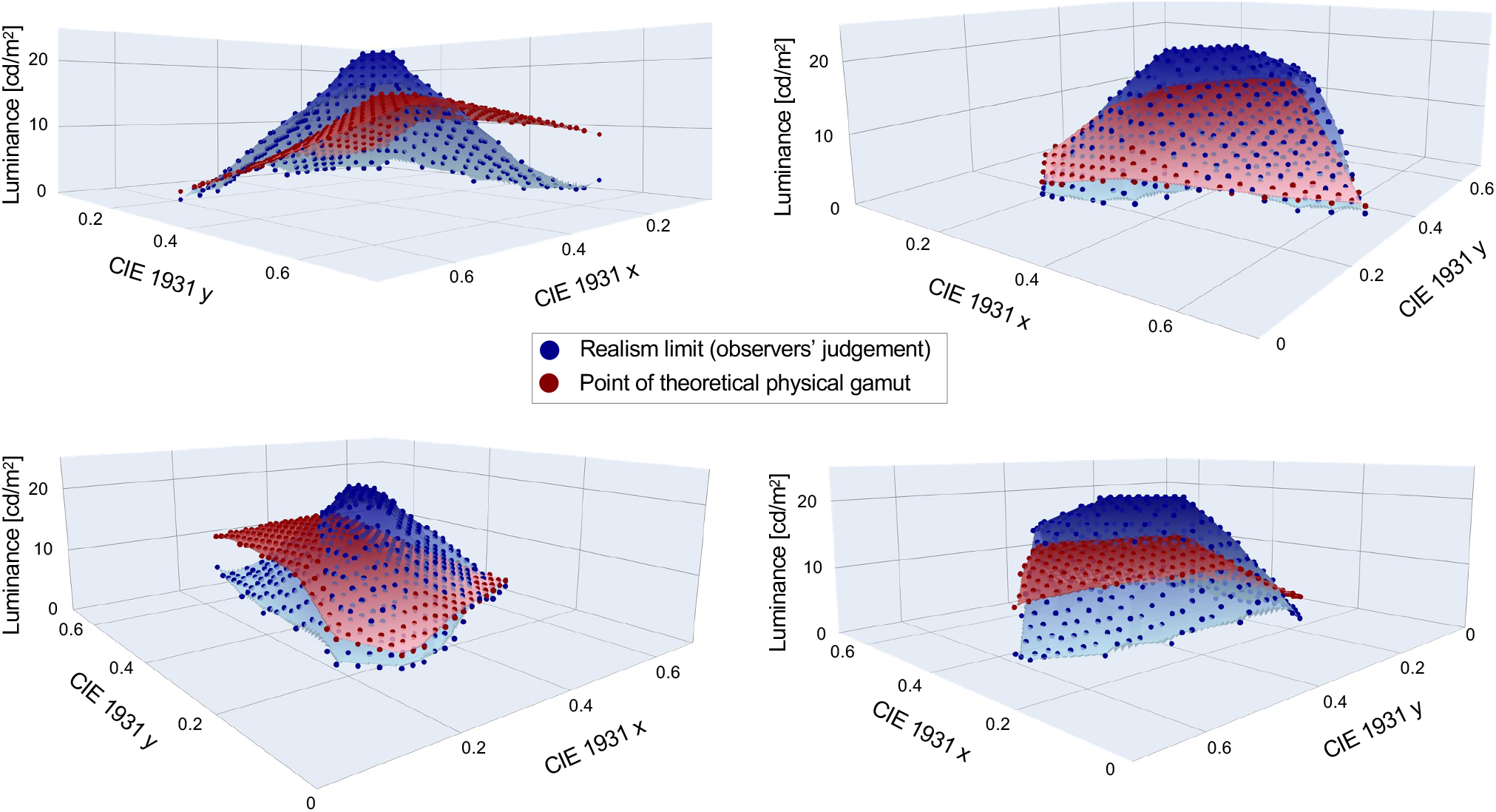
Observers’ judgements of realism and theoretical physical gamut under illuminant 6500 K. The theoretical physical gamut (in red) and the observers’ judgements of realism (in blue) are plotted in the CIE 1931 *xyY* color space for visual comparison. The points represent the actual data, while the surfaces are linear interpolations between these points intended to provide better visualization.

**Figure 2.**
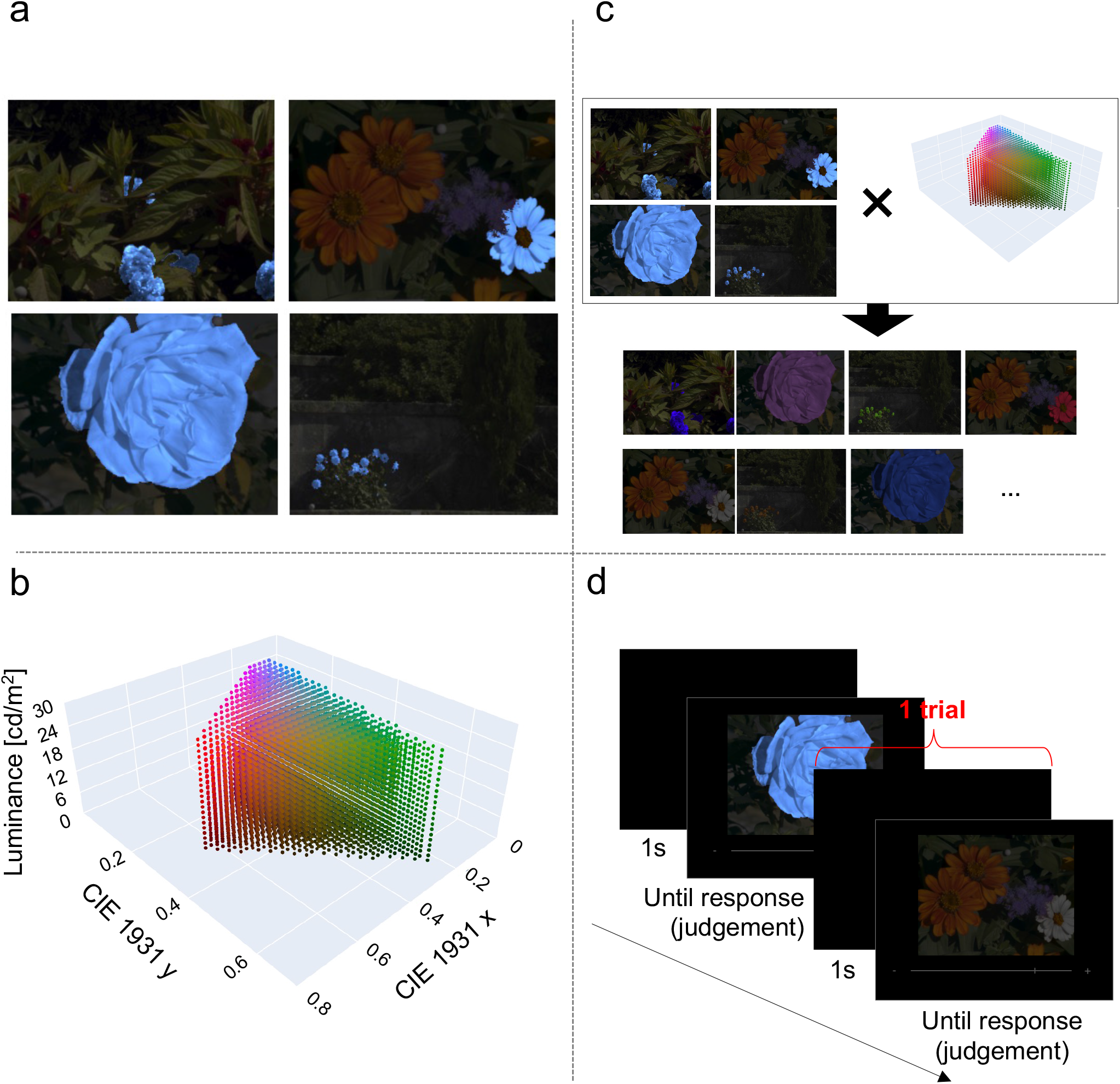
Methods and experimental procedures. Panel **a** represents the natural scenes stimuli used in our experiments; the background and the target are shown, with the target highlighted in blue to clearly illustrate which part of the image was changed throughout the trials. Panel **b** represents the complete set of colors tested and applied to the targets throughout the trials, plotted in the CIE 1931 *xyY* color space. Panel **c** illustrates the stimulus generation procedure, showing how the test color is applied to the target within the stimuli. Panel **d** illustrates the procedure for two experimental trials.

A Pearson correlation analysis was conducted to evaluate the results. The Pearson correlation coefficient between observers’ judgements and the theoretical physical gamut was *r* = 0.63 (*p* < 0.001, 95% CI = [0.54, 0.71]; see Methods for analysis procedure). This result indicates a moderate-to-strong relationship, suggesting that the theoretical physical gamut can predict perceived realism to a certain extent. Consequently, our hypothesis is partially supported, although the model remains imperfect. A visual examination of the point cloud distributions in Fig. 1 and its interactive 3D counterpart Supplementary Fig. S1 online reveals two primary flaws. First, while the overall shapes appear similar, observers’ judgements exhibit a higher vertical magnitude than the theoretical physical gamut along the luminance (*Y*) axis. Second, a notable misalignment is observed in the region associated with low *x* and high *y* coordinates, a range that specifically corresponds to the saturated green area of the CIE 1931 *xy* chromaticity diagram. These two observations are analyzed and discussed in further detail in the following subsections.

### Observers’ judgements exhibit a higher luminance magnitude

While observers’ judgements exhibit a higher luminance magnitude than the theoretical physical gamut, these results are consistent with a similar trend that has been reported in studies investigating luminosity thresholds and physical gamut^13,14^. The authors attribute this phenomenon to the tendency of the human visual system to overestimate the luminous intensity of a scene. Indeed, the physical gamut theory is primarily predicated on the estimation of a scene’s illuminant, from which the visual system derives the physical gamut to subsequently infer perceptual judgements. Consequently, if the scene illuminant is overestimated in intensity, results such as those depicted in Fig. 1 and its interactive 3D counterpart Supplementary Fig. S1 online are to be expected. The degree of illuminant overestimation reported in our results aligns with values observed in these prior studies on luminosity thresholds^13,14^. Furthermore, a similar overestimation of the scene illuminant has been previously observed in color constancy studies^17,24^.

### A misalignment is observed in the green region of the chromaticity diagram

To investigate whether the green region accounts for the observed discrepancies, we performed a leave-one-out (LOO) analysis to identify the data points that most deteriorate the Pearson correlation between the observers’ judgements and the theoretical physical gamut (see Supplementary Methods 3 online for further details). The results of this analysis are presented in Fig. 3. These findings corroborate the trend observed in Fig. 1 and Supplementary Fig. S1 online, confirming that the region corresponding to highly saturated greenish chromaticities is anomalous. After excluding the most detrimental data points (highlighted in orange in Fig. 3b), we recalculated the Pearson correlation coefficient, obtaining *r* = 0.89 (*p* < 0.001, 95% CI = [0.84, 0.92]). This result suggests that the theoretical physical gamut provides a robust framework for predicting perceptual realism for nearly all chromaticities, failing only to account for those in the high-saturation green range.

**Figure 3.**
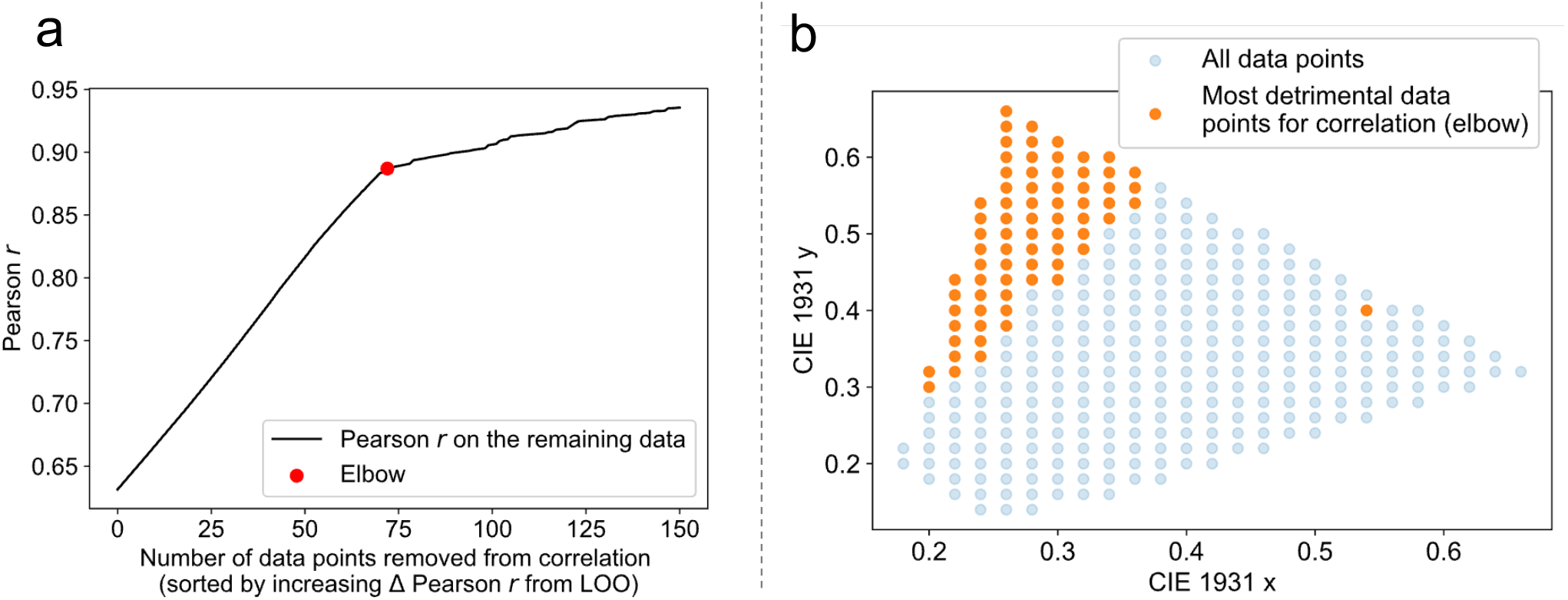
The most detrimental points for the correlation. Panel **a** shows the evolution of the Pearson correlation (vertical axis) as the most detrimental points are removed (horizontal axis). The red dot marks the “elbow” position, which signifies the end of a meaningful improvement in the correlation. Panel **b** plots the most detrimental points revealed by the elbow in Panel **a** on the CIE 1931 *xy* chromaticity diagram. These data pertain to the main experiment conducted under the 6500 K illuminant.

One potential account for this discrepancy could be that such chromaticities are absent from the real-world or exhibit lower maximum luminance levels than those predicted by the mathematical framework of the theoretical physical gamut. Specifically, chlorophyll-rich vegetation absorbs most light, reflecting only a limited amount^25–27^. Under the physical gamut hypothesis, the visual system develops an internal reference via evolutionary exposure to the world. If these saturated green colors do not occur naturally, they remain unlearned, meaning an idealized mathematical model cannot serve as a reliable predictor of their perceived realism. To investigate this hypothesis, we modeled a *real-world physical gamut* by integrating 3,755 empirical reflectance measurements (sourced from the USGS^19^ and ECOSTRESS^20,21^ spectral libraries) into the CIE 1931 *xyY* color space under the experimental illuminant, followed by a final interpolation yielding the gamut’s boundaries (see Methods for further details). The modeling of the real-world physical gamut is displayed in Fig. 4 (and the corresponding interactive 3D counterpart Supplementary Fig. S2 online) and plotted alongside observers’ judgements of realism in Fig. 5 (and the corresponding interactive 3D counterpart Supplementary Fig. S1 online). For an enhanced visualization experience, we recommend consulting the interactive versions online. These results demonstrate that the geometry of the real-world physical gamut aligns remarkably well with that of the observers’ judgements. Figure 6a-c presents the results projected onto the CIE 1931 *xy* chromaticity diagram. This representation provides a top-down view in which the color of the plotted regions corresponds to luminance values normalized for comparability across datasets. The panels respectively illustrate observers’ judgements, the theoretical physical gamut, and the real-world gamut. The area highlighted in red indicates the chromaticities included in the experimental trials. It is immediately apparent that the real-world physical gamut contains no data points within the green region of the CIE 1931 *xy* diagram; in other words, no reflectance spectra corresponding to these chromaticities were present in the USGS or ECOSTRESS spectral libraries. Figure 6d overlays the detrimental data points identified in Fig. 3 onto the real-world physical gamut. This visualization further confirms that the problematic data points correspond to the green region of the chromaticity diagram, where no matching reflectances exist in the spectral databases. This finding lends support to the hypothesis that the theoretical physical gamut cannot effectively predict the perception of realism for saturated green chromaticities, as the human visual system may not have had sufficient prior exposure to such stimuli in the natural environment. However, it should be noted that while this explanation appears plausible, it cannot be fully verified in the current settings. Furthermore, this interpretation relies on a dataset that, despite its considerable breadth, cannot be regarded as exhaustive.

**Figure 4.**
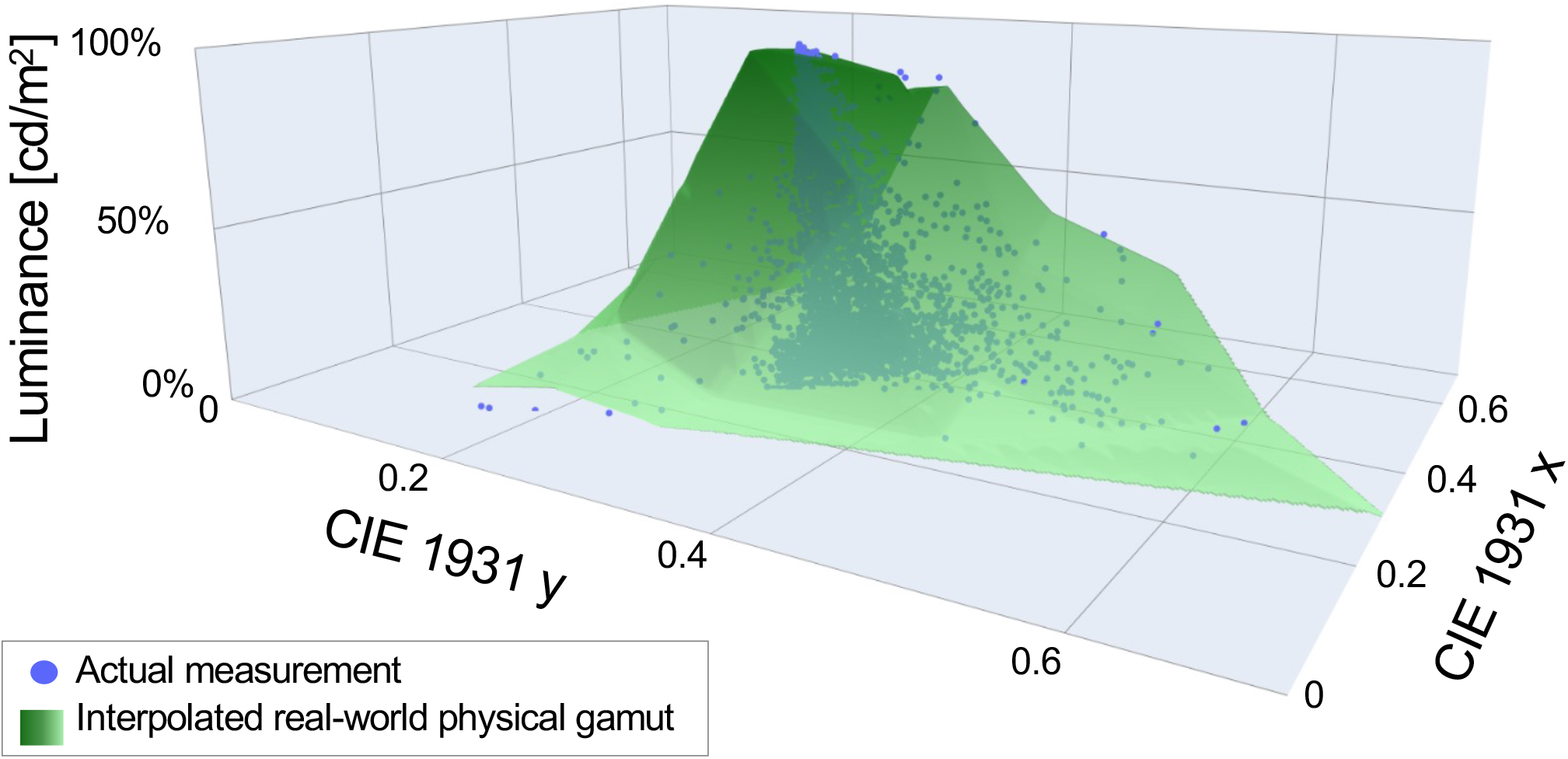
Real-world physical gamut. The real-world physical gamut is represented in the CIE 1931 *xyY* color space. The blue dots show the data obtained from the measurements of USGS^19^ and ECOSTRESS^20,21^ spectral libraries, while the green surface shows the interpolated version of the real-world physical gamut. The luminance axis is represented in relative values because it depends on the intensity of the illuminant (15 cd/m^2^ at 100% of luminance in our experiment).

**Figure 5.**
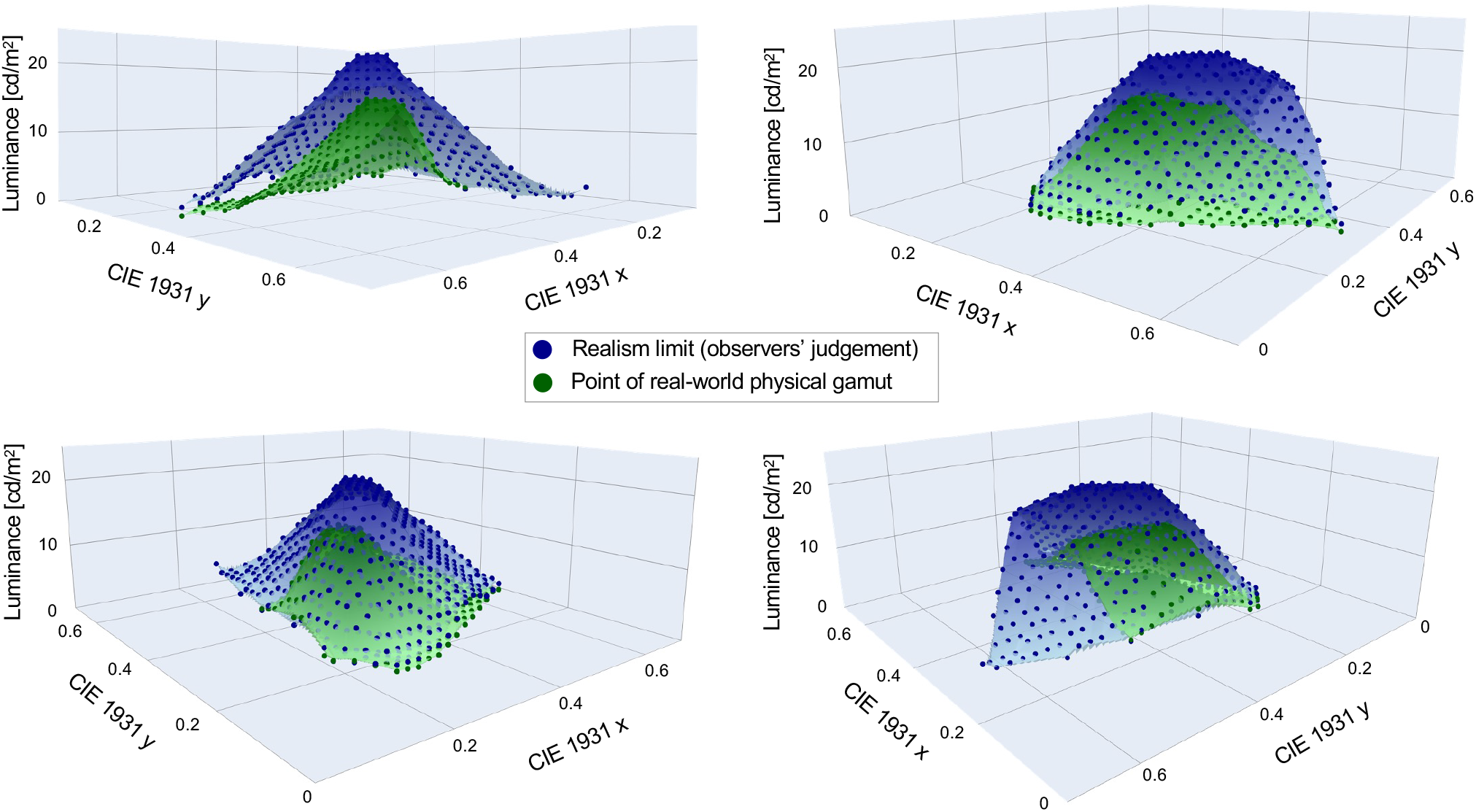
Observers’ judgements of realism and real-world physical gamut under illuminant 6500 K. The real-world physical gamut (in green) and the observers’ judgements of realism (in blue) are plotted in the CIE 1931 *xyY* color space for visual comparison. The points represent the actual data, while the surfaces are linear interpolations between these points intended to provide better visualization.

**Figure 6.**
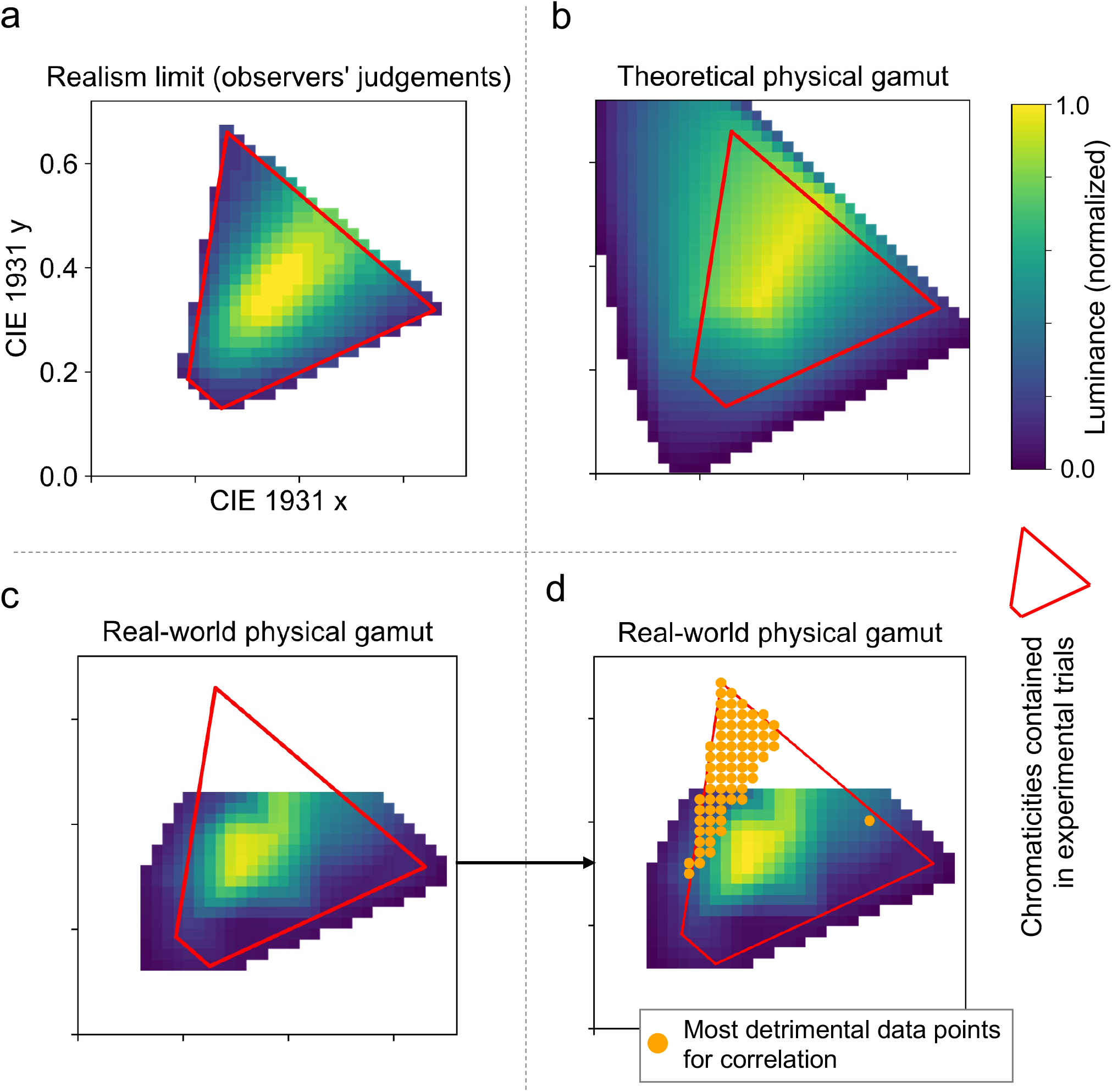
Comparison between theoretical physical gamut and real-world physical gamut. Panel **a** shows the observers’ judgements of realism. Panel **b** shows the theoretical physical gamut. Panel **c** shows the real-world physical gamut. Panel **d** shows the most detrimental points to the correlation (also shown in Fig. 3), plotted on the real-world physical gamut. The area outlined in red in each panel indicates the chromaticities included in the experimental trials. Each panel plots the data on the CIE 1931 *xy* chromaticity diagram, where the color of the plot represents the normalized luminance (*Y*) to provide a more intuitive visual comparison. These data pertain to the main experiment conducted under the 6500 K illuminant.

In addition, we also computed the Pearson correlation coefficient between the interpolated real-world physical gamut and observers’ judgements of realism, following the same procedure as that used for the theoretical physical gamut. This analysis yielded a correlation of *r* = 0.89 (*p* < 0.001, 95% CI = [0.85, 0.92]). Since the interpolated real-world physical gamut contains fewer data points than the observers’ judgements (cf. Fig. 6a and c), we restricted the analysis to the intersection of both datasets, resulting in 263 common points. For these same 263 points, the Pearson correlation between observers’ judgements and the theoretical physical gamut was *r* = 0.80 (*p* < 0.001, 95% CI = [0.73, 0.85]). These findings demonstrate that the real-world physical gamut not only outperforms the theoretical physical gamut but, notably, can serve as a highly effective predictor of realism judgements. This predictive power is particularly striking given that the real-world gamut is derived from a relatively limited number of reflectance measurements. Although outperformed by the real-world model, the theoretical physical gamut nonetheless exhibits a relatively strong correlation with the current dataset. This result suggests that the theoretical physical gamut remains a robust model for both approximating the real-world physical gamut and predicting realism perception, presumably provided that natural objects exist in the real-world for the chromaticities included in the model. Moreover, the strong correlation between the real-world model and realism judgements provides indirect support to the internal reference hypothesis. Indeed, these results tend to indirectly support the assumption that the human visual system might rely on previous empirical observations of the world. Finally, it is reasonable to posit that increasing the number of reflectance measurements and enhancing the physical fidelity of the real-world model would further improve its precision, leading to even more robust correlations and predictive power.

Finally, to assess the generalizability of these findings under different lighting conditions, the experiment was replicated by changing the illuminant from 6500 K to 3000 K. The results were consistent with our previous observations, further supporting the hypothesis that physical gamuts, whether theoretical or empirical, correlate with realism judgements and serve as reliable predictors of the perception of realism. Comprehensive details and results for this supplementary experiment are provided in Supplementary Methods 1 online.

## Discussion

In the present study, we investigated the hypothesis that theoretical physical gamut, derived from optimal colors, can serve as a predictor of realism in natural scenes. To test our hypothesis, we conducted a psychophysical experiment in which observers evaluated the realism of natural scenes depicted in images. Across experimental trials, various colors were applied to a target within the scenes, ensuring that the CIE 1931 *xyY* color space was comprehensively sampled.

The results indicate that the theoretical physical gamut model is highly correlated to realism judgements and can be utilized as an effective predictor of realism perception in natural scenes, provided that the highly saturated green region of the CIE 1931 *xy* chromaticity diagram is excluded. Specifically, the model exhibits a performance deficit within the saturated green region. A plausible explanation for this discrepancy could be that the saturated green chromaticities are presumably non-existent in the natural world. Given that the theoretical framework of physical gamut is predicated on the empirical construction of an internal reference through the observation of real-world, the human visual system would not have been able to internalize or integrate these chromaticities. Consequently, a divergence would occur between observers’ judgements of realism and the theoretical physical gamut, whereas the latter is based on an idealized mathematical definition rather than physical reality. Although this explanation cannot be definitively verified, a supplementary analysis lent it weight. Indeed, we modeled a real-world physical gamut relying on empirical reflectance measurements sourced from spectral libraries and the results exhibit no spectra for the saturated green region of the CIE 1931 *xy* chromaticity diagram. One could argue that even if specific chromaticities are absent from nature, their presence in anthropogenic objects might contribute to the formation of the human visual system’s internal reference. However, we believe that the theoretical framework of physical gamut posits that such internal references would have been shaped by empirical observations over evolutionary timescales rather than the relatively recent modern era or within the span of an individual human lifetime, though this remains speculation that cannot be verified. One might further contend that, since our experimental stimuli consisted of natural targets (flowers), the use of artificial objects might have yielded judgements more closely aligned with the theoretical physical gamut. This possibility remains speculative, as the aforementioned constraint may still apply regardless of the stimulus category. A comparative investigation between natural and anthropogenic targets could serve as the subject of future research. It is also worth noting that this analysis is predicated on the assumption that the spectral libraries utilized (USGS^19^ and ECOSTRESS^20,21^) contain datasets sufficiently large to derive response trends. Although these libraries are objectively extensive and encompass a vast population of materials and objects making up the real-world (including man-made objects), we cannot draw strong or definitive conclusions. Consequently, we draw attention to the exploratory and non-definitive nature of this supplementary analysis. Nevertheless, assembling a database more extensive than those utilized in the present study remains a significant challenge; such an endeavor could also serve as the subject of future work.

Moreover, the aforementioned real-world physical gamut was found to be notably strongly correlated with observers’ judgements, serving as a potential very effective model for predicting the perception of realism. Lastly, a subsequent experiment conducted under a different daylight illuminant suggests that these findings are not limited to a single condition but may generalize across different illuminants, thereby strengthening the proposed theory.

This study provides further insight into how realism is perceived in natural scenes and which mechanisms appear to contribute to this process. It may prove beneficial for future research on these topics or for other studies concerning color preference, echoing the research fields discussed in our introduction. Furthermore, it can be applied to address more tangible challenges within the engineering community, such as realistic graphical rendering of virtual objects or environments. More specifically, these findings could be utilized in Augmented Reality (AR) or Mixed Reality (MR). Indeed, these technologies aim to integrate virtual objects into the physical world with the ultimate objective of creating a seamless illusion where it is impossible to distinguish the virtual from the real. One of the primary challenges in achieving this goal is the integration of virtual objects while faithfully adhering to illumination constraints, ensuring that colors and luminances remain plausible within a given real-world context. The physical gamut models (both the theoretical and real-world variants) presented in this study help define the boundaries that virtual objects must respect to be perceived as realistic within Extended Reality (XR) experiences.

## Methods

### Apparatus

The experiments were conducted in a darkroom. Visual stimuli were presented on a 25-inch Sony PVM-A250 OLED monitor (Sony, Japan) featuring a spatial resolution of 1920 × 1080 pixels, a 10-bit color depth per channel, and a refresh rate of 60 Hz. To ensure accurate reproduction of luminance and chromaticity, the display was rigorously calibrated using a spectroradiometer (Specbos1211-2, JETI Technische Instrumente GmbH, Germany) and a colorimeter (ColorCAL II, Cambridge Research Systems, UK). The hardware setup consisted of the monitor interfaced with a MacBook Air (Apple M2, macOS Sequoia 15.2; Apple, USA), while experimental control was managed through custom scripts developed in PsychoPy (version 2024.2.4). Observers were positioned at a viewing distance of 40 cm, with head stability maintained by a chinrest. Judgements responses were registered using a standard computer mouse.

### Observers

A total of 22 participants (mean age = 23.86, SD = 3.05; 2 females, 20 males) took part in the research. The investigation was carried out from May 13, 2025, to June 6, 2025, with observers recruited through onsite solicitation at the university during this interval. All individuals were undergraduate or graduate students affiliated with the Institute of Science Tokyo. Every participant successfully passed the Ishihara color vision assessment and exhibited normal or corrected-to-normal visual acuity. The experimental design adhered to the Declaration of Helsinki and was approved by the Ethical Review Committee of the Institute of Science Tokyo. Informed written consent was obtained from all observers after the experimental protocols were thoroughly explained.

### Color computation

Within the field of color science, the CIE 1931 *xyY* color space offers a conventional framework for the quantification of color as perceived by the human visual system. The calculation of colors within this system necessitates three fundamental elements^28^: the spectral reflectance of the surface, the spectral power distribution (SPD) of the light source, and the CIE 1931 color matching functions (CMFs). These functions characterize the spectral sensitivity of a standard human observer, specifically the 2^◦^ Standard Observer utilized in this research. Spectral reflectance signifies the proportion of light reflected by a surface at specific wavelengths, whereas the SPD of the illuminant specifies the intensity of incident light across the spectrum. The spectral composition of the reflected light is determined by calculating the product of the reflectance and the illuminant at each wavelength. This resulting product is subsequently integrated over the visible spectrum and weighted by the CMFs to produce the final color represented as tristimulus values (*X,Y, Z*), from which the chromaticity coordinates (*x, y*) and luminance *Y* are derived. Hyperspectral imaging provides comprehensive spectral reflectance data for every pixel, allowing for the precise determination of color for each pixel under any specified illuminant. Further technical details can be found in established literature^28,29^. The present study employed this methodology to generate the experimental stimuli and colors.

### Optimal colors and theoretical physical gamut

Optimal colors are theoretically defined as surfaces with spectral reflectance functions consisting strictly of 0% and 100% values, characterized by a maximum of two abrupt transitions across the wavelength spectrum^13,15–18^. In the real-world, however, physical constraints prevent any material from achieving more than 100% reflectance at any given wavelength. Consequently, optimal colors represent the theoretical upper luminance limit for any specific chromaticity; no naturally occurring surface can exceed the luminance of these theoretical distributions. Therefore, by calculating the exhaustive set of optimal reflectance functions and spectrally multiplying them by a specific light source, one can determine the theoretical physical gamut permissible under that illuminant. For this study, we generated optimal colors within the CIE 1931 color space following the guidelines of established literature^13,17,18^. Specifically, we derived approximately 90,000 optimal spectral reflectance functions by scanning all possible combinations of wavelengths that respect the optimal colors criteria within the 400–700 nm range at a 1 nm sampling interval. Each resulting spectrum was multiplied by the spectral power distribution of a 6500 K Planckian radiator and integrated into the CIE 1931 color space following the aforementioned methodology. The resulting optimal colors set defines the theoretical physical gamut for the 6500 K illuminant condition used in our experiment. The generated gamut is illustrated in Fig. 1 and its corresponding 3D interactive visualization Supplementary Fig. S1 online. It should be noted that while the initial physical gamut is calculated in normalized luminance, the luminance values presented in Fig. 1 and Supplementary Fig. S1 online have been scaled by the scene’s illuminant intensity (fixed at 15 cd/m^2^ in our experiments) to reflect absolute luminance levels.

The conceptual and practical foundations of optimal colors (including the computation of color coordinates) are extensively documented in literature. Accordingly, this manuscript provides the essential information required to contextualize the present study while omitting redundant technical derivations. For a comprehensive overview of the underlying computational procedures, the reader is referred to the relevant references^13,17,18^.

### Real-world physical gamut

A real-world physical gamut was constructed based on empirical reflectance measurements. Approximately 6,000 reflectance spectra were aggregated from two publicly available repositories: the USGS^19^ and ECOSTRESS^20,21^ spectral libraries. These datasets encompass a comprehensive range of materials found in natural environments and terrestrial landscapes, including man-mades materials and objects. The data were curated and pre-processed to suit the specific requirements of this study. Specifically, the following categories were retained: man-made materials, coatings, liquids, minerals, organic compounds, soils, and mixtures. Conversely, lunar surface measurements and samples lacking data within the visible spectrum (spanning the wavelength range of 400 nm to 700 nm) were excluded. Utilizing the metadata of the measuring instruments provided by the source databases, reflectance values were converted to percentages. Samples exhibiting reflectance levels exceeding 100%, which may indicate phosphorescent or fluorescent properties, were subsequently omitted. This filtering process yielded a final dataset of 3,755 measurements. These reflectance spectra were then integrated into the CIE 1931 *xyY* color space, following the same methodology previously described. Finally, a three-dimensional bin interpolation (without extrapolation) was performed to characterize the volume and derive an enveloping three-dimensional hull for the measurements. The *xy* chromaticity plane was partitioned into a 12 *×* 12 grid. Within each bin, the data point with the highest luminance was selected and re-assigned to the *xy* coordinates of the bin center. Linear interpolation was then applied between these peak points. This procedure allowed for the definition of the *real-world physical gamut*, as referred to throughout this article. The resulting gamut and the former reflectance measurements are illustrated in Fig. 4 and its corresponding 3D interactive visualization Supplementary Fig. S2 online.

### Stimuli

The objective of this experiment was to evaluate observers’ realism judgements across four distinct natural scenes. In each scene, a specific target region was manipulated across trials by varying its color (chromaticity and luminance) to span the available CIE 1931 *xyY* color space permitted by the monitor’s gamut.

Four natural scenes were selected from a publicly available dataset of hyperspectral images^30^ and utilized in previous color vision studies^29,31^. These base images are referred to as *backgrounds* throughout this study. Within each background, a target object was defined for color manipulation. Flowers were chosen as targets because their natural diversity allows them to exhibit a wide range of colors without introducing cognitive bias or appearing inherently unrealistic to the observer. Figure 2a illustrates the four backgrounds and their respective targets (highlighted in bright blue for better visualization).

To manipulate the targets, we defined a comprehensive set of test colors by sampling the CIE 1931 *xyY* color space. The chromaticity coordinates (*x, y*) were sampled in increments of 0.02 from 0.0 to 1.0 on both axes, while luminance (*Y*) was sampled in 20 discrete steps of 1.15 cd/m^2^ from 2 cd/m^2^ to 25 cd/m^2^. We retained only the colors reproducible accurately within the gamut of the monitor used in the study, resulting in a total of 7,587 test colors. Figure 2b visualizes this distribution of test colors within the CIE 1931 *xyY* chromaticity diagram.

The final experimental set comprised 30,348 images (7,587 colors applied to four backgrounds). Backgrounds were initially generated by multiplying the hyperspectral data by the spectral power distribution of a 6500 K Planckian radiator. The illuminant intensity was scaled such that a perfectly reflecting surface within the image (100% reflectance across all wavelengths) would yield a luminance of 15 cd/m^2^. A relatively low background luminance was intentionally selected to rigorously test the physical gamut hypothesis. As the physical gamut concept is fundamentally defined by luminance limits, a low-luminance background ensures that observers can clearly perceive luminance contrasts when the target luminance is high. Since human contrast sensitivity decreases at higher luminance levels, a high-luminance background would likely reduce the precision of realism judgements by saturating the visual response. The spectral product was then integrated using the CMFs (CIE 2^◦^ Standard Observer) to derive CIE 1931 (*x, y,Y*) coordinates. Subsequently, a mask was employed to isolate and manipulate target pixels. While chromaticity (*x, y*) was applied uniformly to the target region, the original luminance structure was preserved to maintain the image’s texture and spatial details. This was achieved by normalizing the original target luminance to a [0, 1] range and subsequently multiplying it by the target luminance value (*Y*) from the test set. This procedure is illustrated in Fig. 2c. Finally, the images were converted from CIE 1931 *xyY* space to RGB space based on the monitor calibration procedure described in the next section.

### Color calibration

To maintain rigorous control over the presented stimuli, the monitor was calibrated according to established protocols common in color science research^32,33^. Within this framework, the display’s behavior is characterized by the spectral power distributions of the three primary colors and the luminance-response (gamma) functions for each color channel. These parameters allow for the prediction of CIE 1931 *XYZ* output based on digital RGB inputs; conversely, they facilitate the derivation of the specific RGB values needed to achieve a desired CIE 1931 *XYZ* (or *xyY*) tristimulus. Primary spectra were recorded at maximum intensity using a spectroradiometer, while the gamma functions were characterized via a colorimeter (technical details for both instruments are provided in the Apparatus section). Although these methodologies were originally developed for older hardware, the same principles are widely applied to modern additive RGB displays, provided their primaries demonstrate the spectral stability and channel independence required for this calibration model. Therefore, in alignment with standard research practices, this procedure was utilized for our display and experimental software. To ensure long-term accuracy, calibration was updated on a weekly basis. Furthermore, because OLED technology requires thermal equilibrium, the monitor was warmed up for 30 minutes before any measurement or experimental session. During this period, dynamic video content was displayed to maintain pixel activity and mitigate the risk of image retention. This rigorous approach ensures that any CIE 1931 *XYZ* (or *xyY*) tristimulus within the display’s color space can be reproduced at the pixel level with a margin of error within one unit of the *CIEDE*2000 color-difference formula. To prevent signal clipping, all experimental stimuli were selected to remain within the achievable gamut of the display.

### Experimental procedure

Each experimental trial consisted of a subjective realism assessment. Observers were instructed to evaluate the overall realism of a stimulus on a scale ranging from 0 to 10. The stimulus was presented in the center of the display. The stimuli subtended a visual angle of 13.26° horizontally and 10.0° vertically. To ensure high contrast and a true black level, stimuli were presented on an OLED monitor with a background luminance of 0 cd/m^2^. Below the stimulus, an achromatic horizontal scaling bar (luminance of 8 cd/m^2^) was displayed. Observers adjusted a cursor along this bar using horizontal mouse movements and confirmed their judgement with a right-click. The scale featured achromatic “+” and “-” symbols (luminance of 8 cd/m^2^) at the right and left poles, respectively, to indicate the direction of the rating. While the bar contained no numerical markers other than the cursor, observers were briefed that the range represented a continuum from 0 (“not realistic at all”) to 10 (“completely realistic”), with 5 serving as the qualitative threshold between unrealistic and realistic. Participants were explicitly instructed to judge the image holistically. No time constraints were imposed on the response period, and a 1-second black-screen inter-trial interval (ITI) was maintained. Figure 2d illustrates the sequence of two representative trials.

Each experimental session began with a 2-minute dark adaptation period featuring a black screen. The dark adaptation was followed by 40 practice trials designed to familiarize observers with the full range of chromaticities and luminances used in the study. These practice stimuli were consistent across all observers and sessions, though presented in a randomized order. The practice set comprised 10 distinct chromaticities spanning the entire display gamut, each presented at four luminance levels: 2 cd/m^2^ (minimum stimulus luminance), 6.6 cd/m^2^ (intermediate), 15 cd/m^2^ (maximum physical luminance under the reference illumination), and 25 cd/m^2^ (maximum stimulus luminance). Beyond familiarization, these trials served to allow chromatic adaptation to the experimental illuminant. The practice phase lasted approximately 2 minutes on average, exceeding the standard 1-minute adaptation period typically observed in color vision protocols. Data collected during this phase were excluded from any analysis. Upon completion, a brief achromatic message (luminance of 8 cd/m^2^) signaled the transition to the formal experimental trials.

The total stimulus set of 30,348 images was distributed across 22 observers. Each observer completed 1,379 or 1,380 trials, partitioned into five sessions of 275 or 276 trials each. All five sessions were conducted within a single day, with observers permitted self-paced breaks between sessions to mitigate fatigue. As detailed in the Stimuli section, the 30,348 trials resulted from applying 7,587 test colors to the targets within four distinct background images. Each unique color-background combination was assigned to a single observer and was not repeated for that individual. However, test colors were indirectly sampled four times across the study, as each was paired with the four different backgrounds. Due to the randomized assignment protocol, the four background variations for a given test color were evaluated by different observers.

### Analysis

The primary objective of this study was to test the hypothesis that theoretical physical gamuts predict realism judgements across the entirety of the color space, rather than within a restricted subset of colors. To rigorously evaluate the extent to which these gamuts account for the perception of realism, it is necessary to sample the color space at the highest possible resolution. Because this approach requires an extensive set of color stimuli, thereby significantly increasing the number of trials and the overall experimental duration, we implemented a pseudo big-scale strategy. This involved recruiting a larger number of observers than the standard in color vision research to distribute a high volume of trials, ensuring dense coverage of the CIE *xyY* space. Our goal was to elicit judgements for a vast number of finely spaced colors, replicated across four distinct images. This methodological framework provides a substantial dataset from which we can derive robust trends within the three-dimensional color space. Specifically, by testing a high density of colors and repeating this coverage across four different backgrounds, we compensate for the lack of individual repetitions for each color-background combination. This allows for the extraction of stable, global trends across the color space rather than relying on precise judgements for isolated colors. These trends are subsequently compared to the theoretical physical gamut to verify whether the latter serves as a reliable predictor of perceived realism.

To robustly capture the underlying trends in observers’ judgements and mitigate the noise inherent in the large-scale experimental design, a multi-stage smoothing procedure was applied to the data. Initially, the judgements from 22 observers, encompassing a total of 30,348 trials, were averaged for each CIE 1931 (*x, y,Y*) coordinates. This averaging was performed across all background conditions, as each color was presented once per background. Subsequently, the averaged judgements underwent smoothing via locally weighted linear regression utilizing a Gaussian kernel within the three-dimensional CIE 1931 *xyY* color space. For each data point, a local affine model was fitted to the judgements, with weights determined by an anisotropic Gaussian kernel that assigned greater influence to points in closer proximity in both chromaticity (*x, y*) and luminance (*Y*). Given the disparate sampling resolutions along the chromaticity axes (steps of 0.02) and the luminance axis (steps of 1.15 cd/m^2^), an ellipsoidal rather than a spherical kernel was employed to ensure appropriate spatial weighting. This anisotropy was defined through the bandwidth parameters *h*_*xy*_ and *h*_*Y*_, which were set to twice the respective sampling resolutions, resulting in *h*_*xy*_ = 0.04 and *h*_*Y*_ = 2.3. The model was evaluated at every data point to obtain locally averaged estimates, and these smoothed judgements were used to replace the original observers’ responses. A comprehensive mathematical formalization of this smoothing procedure is provided in Supplementary Methods 2 online.

The smoothing of observers’ ratings resulted in a point cloud within the CIE 1931 *xyY* color space, where each point was associated with a fourth-dimensional value representing the realism judgement. To define the boundary of perceived realism, we first filtered the dataset to include only points with a realism score greater than or equal to 5 (the predefined threshold between realistic and unrealistic perceptions communicated to observers during experimental sessions). Following this filtering procedure, all (*x, y*) chromaticity coordinates remained represented at least once within the dataset. For each chromaticity coordinate (*x, y*), we retained only the data point corresponding to the maximum luminance *Y*. This transformation converted the initial volumetric point cloud into a hollow shell consisting of 345 three-dimensional points, representing the upper limits of perceived realism across the sampled color space. This surface is visualized in Fig. 1 and its interactive 3D counterpart Supplementary Fig. S1 online.

The theoretical physical gamut is a model defining the maximum luminance a chromaticity can achieve under a specific illuminant. In the three-dimensional CIE 1931 *xyY* space, this is represented as a hollow 3D cone defining the physical boundaries of color: each chromaticity (*x, y*) is paired with a maximum luminance value *Y*. Our hypothesis posits this model as a potential predictor of the threshold between realistic and unrealistic stimuli, where the realism boundary corresponds to the maximum physical luminance defined by the gamut. Although the theoretical physical gamut was initially generated for the entire CIE 1931 space, our experimental trials were constrained by the color gamut of the monitor. Consequently, for the analyses and visualizations in Fig. 1 and Supplementary Fig. S1 online, the theoretical gamut was filtered to include only the chromaticities present in the experimental trials.

The final datasets comprised two sets of 345 points, each forming a 3D surface in the CIE 1931 *xyY* color space: the empirical observers’ judgements and the theoretical physical gamut. To test our hypothesis, we calculated the Pearson correlation coefficient between these two datasets. The 345 empirical judgements and their corresponding theoretical predictions were organized into two one-dimensional vectors, sorted by chromaticity (*x, y*), to allow for a point-by-point correlation.

To evaluate the statistical significance of the observed correlations, we applied a two-tailed, non-parametric bias-corrected and accelerated (BCa) bootstrapping method with 10,000 resamples at the observer level and a significance level of 5%^34^. For each bootstrap sample, observers were resampled with replacement (N = 22). For each iteration, a new sample was drawn, and the corresponding observers’ judgements were retrieved. These data were averaged for each CIE (*x, y,Y*) coordinate across backgrounds, ensuring that observers selected multiple times during resampling were first averaged at the individual observer level before being aggregated across background conditions. The resulting averaged dataset was then smoothed using the aforementioned local linear regression model, from which the realism limits were extracted. Finally, the Pearson correlation coefficient *r* between these resampled limits of realism and the theoretical physical gamut was calculated. This iterative process yielded a distribution of 10,000 Pearson *r* values, from which the *p*-value and the 95% confidence intervals were derived to determine the robustness of the results.

## Data availability

All data generated or analysed during the current study are available from the corresponding author on reasonable request.

## Additional Information: Competing interests

The authors declare no competing interests.

## Acknowledgements

This study was supported by JST SPRING, Japan Grant Number JPMJSP2180 to KD, and JSPS KAKENHI Grant Number 23K28174 and 21KK0203 to TN.

## Author contributions statement

KD and TN designed the experiment and interpreted the results. KD conducted the experiments, analyzed the results, and wrote the manuscript. All authors reviewed and approved the final version of the manuscript.

