## Supplementary Methods 1, 2, and 3 for "The perception of realism is correlated with the concept of physical gamut"

### 1 Supplementary Information

#### 1.1 Supplementary Methods

##### 1.1.1 Supplementary Methods 1: Replication of experiment under the illuminant 3000 K

The main experiment conducted under a 6500 Kelvin (K) illuminant was replicated using an alternative light source to evaluate the generalizability of the findings. Since physical gamut theory is predicated on the observer's internal estimation of the illuminant, it is noteworthy to validate the results under a secondary lighting condition. This replication allows for an assessment of whether perceptual judgements adapt accordingly and whether the high degree of correlation remains consistent. For this purpose, a 3000 K illuminant was selected, representing a significantly warmer, orangish spectral distribution.

We generated the theoretical physical gamut, the stimuli, and the real-world physical gamut following the same protocol as the main experiment, while substituting the illuminant during spectral integration with the spectral power distributions of Planckian radiators at 3000 K.

The experimental design and procedure mirrored those of the main experiment, subject to several specific modifications. Stimulus generation was conducted using a step size of 0.04 within the CIE 1931 $(x, y)$ chromaticity plane, with luminance levels $(Y)$ spanning 2 to 25 cd/m$^2$ across 15 increments of 1.5 cd/m$^2$ (the last increment being rounded to 25 cd/m$^2$). To account for the reduced data resolution in comparison to the main experiment, the ellipse radii utilized for smoothing observers' scalings (see Supplementary Methods 2) were increased to 0.08 for the $(x, y)$ plane and 3.0 for the luminance $(Y)$ axis. These adjustments resulted in a total of 5,256 trials, a reduction from the 30,348 trials used in the primary study. Seven observers participated, each completing two sessions consisting of 375 or 376 trials, resulting in a total of 750 or 751 trials per participant.

Figure 1 in the present material and its interactive 3D counterpart Supplementary Fig. S3 online illustrate the observers' judgements results and the theoretical physical gamut. Similarly, Fig. 2 in the present material and the associated interactive version Supplementary Fig. S3 online present the observers' judgements compared against the real-world physical gamut.

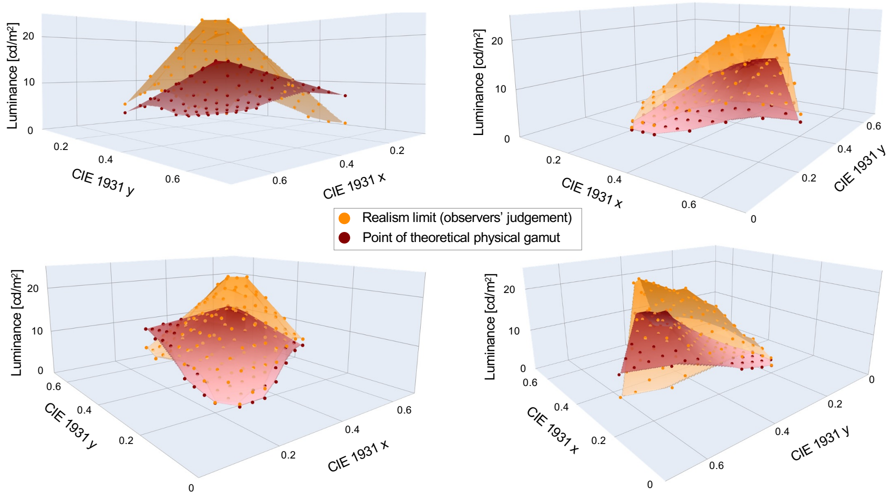


Figure 1: Observers' judgements of realism and theoretical physical gamut under illuminant 3000 K. The theoretical physical gamut (in red) and the observers' judgements of realism (in orange) are plotted in the CIE 1931 $xyY$ color space for visual comparison. The points represent the actual data, while the surfaces are linear interpolations between these points intended to provide better visualization.

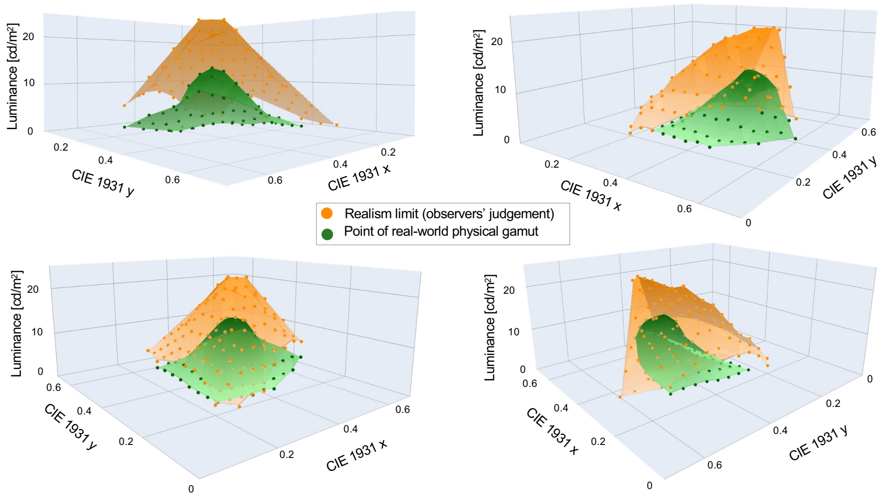


Figure 2: Observers' judgements of realism and real-world physical gamut under illuminant 3000 K. The real-world physical gamut (in green) and the observers' judgements of realism (in orange) are plotted in the CIE 1931 $xyY$ color space for visual comparison. The points represent the actual data, while the surfaces are linear interpolations between these points intended to provide better visualization.

Following the analytical protocol established in the primary experiment, we calculated Pearson correlation coefficients to quantify the relationship between observers' judgements and the respective gamuts. The correlation between observers' judgements and the theoretical physical gamut was $r = 0.68$ ($p < 0.001$, 95% CI = [0.62, 0.72]). A stronger correlation was observed between observers' judgements and the real-world physical gamut, yielding $r = 0.82$ ($p < 0.001$, 95% CI = [0.80, 0.86]). Furthermore, when evaluating the theoretical physical gamut using only the data points corresponding to the real-world comparison, the calculation of the correlation yielded $r = 0.64$ ($p < 0.001$, 95% CI = [0.60, 0.74]).

In conclusion, the findings and trends observed under the 3000 K illuminant are highly consistent with those recorded under the 6500 K condition. This alignment provides further empirical support for our primary hypothesis and suggests that the observed phenomena are robust across varying illuminants. These results indicate that our main findings seem to generalize to other lighting conditions, remaining valid even when the spectral power distribution of the light source is significantly altered. In accordance with the physical gamut theory, observers appear capable of accurately identifying the illuminant and accessing a corresponding internal representation to infer their perception and subsequent judgements.

##### 1.1.2 Supplementary Methods 2: Smoothing of observers' judgements

This supporting information describes the mathematical explanation of how the observers' judgements were modeled and smoothed.

To smooth the realism judgements provided by observers, we employ a locally weighted linear regression model across the CIE 1931 $xyY$ color space. For any target point $\mathbf{q} = [x_q, y_q, Y_q]$, the estimated realism score $\hat{j}$ is obtained by fitting a local linear model that prioritizes neighboring observations through a Gaussian weighting kernel.

The distance $d_i$ between a target point $\mathbf{q}$ and an observation $\mathbf{p}_i = [x_i, y_i, Y_i]$ is defined by a normalized ellipsoidal metric:

$$d_i^2 = \frac{(x_i - x_q)^2 + (y_i - y_q)^2}{h_{xy}^2} + \frac{(Y_i - Y_q)^2}{h_Y^2} \tag{1}$$

The weight $w_i$ assigned to each observation is then calculated using a radial basis function:

$$w_i = \exp\left(-\frac{1}{2}d_i^2\right) \tag{2}$$

The local linear model parameters $\boldsymbol{\beta} = [\beta_0, \beta_1, \beta_2, \beta_3]^T$ are determined by minimizing the weighted least squares objective function:

$$\min_{\boldsymbol{\beta}} \sum_i w_i \left(j_i - (\beta_0 + \beta_1 x_i + \beta_2 y_i + \beta_3 Y_i)\right)^2 \tag{3}$$

Finally, the smoothed realism judgement $\hat{j}$ at point $\mathbf{q}$ is given by the prediction of the fitted local model:

$$\hat{j} = \beta_0 + \beta_1 x_q + \beta_2 y_q + \beta_3 Y_q \tag{4}$$

The parameters of the model are defined as follows:

- $xyY$: Spatial coordinates in the CIE 1931 color space ($x, y$ for chromaticity and $Y$ for luminance);
- $j$: Raw realism judgements collected from observers;
- $h_{xy}$: Smoothing bandwidth (radius) for the chromaticity axes ($x$ and $y$; value fixed at 0.04 for the main experiment under the illuminant 6500 K and 0.08 for the replication under the illuminant 3000 K, according to the logic detailed in the main article and in Supplementary Methods 1);
- $h_Y$: Smoothing bandwidth (radius) for the luminance axis ($Y$; value fixed at 2.3 for the main experiment under the illuminant 6500 K and 3 for the replication under the illuminant 3000 K, according to the logic detailed in the main article and in Supplementary Methods 1).

##### 1.1.3 Supplementary Methods 3: Leave-One-Out (LOO) analysis

This section details the leave-one-out (LOO) analysis methodology employed to identify the data points that most deteriorate the Pearson correlation between the observers' judgements and the theoretical physical gamut in the main article.

First, we established a baseline Pearson correlation coefficient, $r_{\text{all}}$, using the complete set of chromaticities $(x, y)$. For each judgement $i$, the correlation was recomputed upon its exclusion, denoted as $r_{-i}$. We defined the LOO influence of a specific point $i$ as: $\Delta_i = r_{\text{all}} - r_{-i}$. Judgements characterized by more negative $\Delta_i$ values were those whose removal yields the greatest improvement in correlation; consequently, these points were identified as the most detrimental to the correlation. To characterize the improvement in correlation as a function of data exclusion, we ranked all points by $\Delta_i$ in ascending order (from most to least detrimental). We then defined $r(m)$ as the Pearson correlation coefficient calculated after the exclusion of the top $m$ most influential points. The resulting $r(m)$ curve summarizes the trade-off between improving statistical agreement and preserving dataset coverage (see Fig. 3 in the main article).

To determine a principled threshold for data removal, we applied an elbow criterion to the $r(m)$ curve. Heuristically, the elbow represents the transition point toward diminishing marginal returns: prior to this point, the correlation increases sharply as systematic outliers are removed; beyond it, further exclusions merely simplify the dataset without substantially enhancing agreement. We objectively identified this point using the maximum distance to the chord method. A linear segment (chord) was drawn between the endpoints of the $r(m)$ curve over the evaluated range, and the optimal truncation level $m^*$ was defined as the value of $m$ where the perpendicular distance between the curve and the chord was maximized.

Crucially, the objective of this procedure was not to artificially optimize correlation through arbitrary data pruning, but rather to diagnose systematic model failures. Since each observation is associated with a specific chromaticity coordinate $(x, y)$, mapping the $m^*$ excluded points back onto the chromaticity plane facilitates the spatial localization of model discrepancies. If these influential points cluster within specific color regions (e.g., near the green primary), it provides the suggestion that the observed mismatch might be structured. Such clustering can indicate region-specific limitations of the theoretical model rather than noise.
